# Evaluating the effect of badger culling on TB incidence in cattle: a critique of Langton et al. 2022

**DOI:** 10.1101/2025.07.10.664183

**Authors:** Andrew Robertson

## Abstract

The European Badger (Meles meles) is a wildlife host for *Mycobacterium bovis*, the causative agent of bovine tuberculosis (TB) in cattle. In England, large scale badger culling has been implemented since 2013 as part of a licenced Badger Control Policy (BCP). While several studies have shown an association between the BCP and disease benefits in cattle, a study by Langton et al. (2022) failed to identify any evidence of such benefits, raising questions about the effectiveness of badger culling as a tool for disease control. In this study I summarise the primary methodology used in the Langton et al. study and identify two significant issues with the approach used, which compares cattle TB incidence in culled areas of SW England to land without culling. Firstly, their analysis groups data together from cull areas into a combined culled vs unculled treatment, regardless of the time that culling has been applied. Secondly, there is evidence of a clear cull area selection bias, such that land enrolled into culling during the period analysed had a significantly higher baseline TB incidence than areas not culling. I then conducted a Monte Carlo analysis using simulated data under scenarios with differing cull effects, and matching the degree of grouping, selection bias and statistical methods used by Langton et al. (2022). The results indicate that even in scenarios with a relatively rapid and large effect of culling seeded into the simulation there is a significant probability that their analyses will fail to find an effect and may even wrongly conclude that culling is associated with an increase in TB. These results potentially explain the discrepancy between Langton et al. (2022) and other studies evaluating badger culling and highlight the importance of rigorous statistical analysis that accounts for temporal and spatial patterns in disease control data.

## Introduction

Bovine TB, caused by *Mycobacterium bovis* is a globally significant disease, with impacts on cattle, humans and wildlife (Palmer et al. 2012). The European badger acts as a wildlife host for *M. bovis* in the UK, Ireland and parts of mainland Europe, and a broad range of evidence indicates that badgers can act as a source of infection in cattle (Little et al. 1982, Bourne et al. 2007, Donnelly and Nouvellet 2013, Crispell et al. 2019, van Tonder et al. 2021). Large scale field trials have demonstrated that badger culling is associated with reductions in cattle TB incidence in England (Donnelly et al. 2007) and Ireland (Griffin et al. 2005). However, the Randomised Badger Culling Trial (RBCT) conducted in England from 1998-2005 demonstrated that the outcomes of badger culling are not simple or easy to predict and have the potential to both reduce or increase TB incidence in cattle, depending on the scale and approach used (Donnelly et al. 2006, Bourne et al. 2007, Donnelly et al. 2007).

Badgers are a protected native species in the UK and culling is a controversial subject, particularly among some stakeholders and members of the public (Enticott 2015). The results of the RBCT (Donnelly et al. 2007) are also disputed by some studies which question the robustness of the statistical methods used (Torgerson et al. 2024, Torgerson et al. 2025), although other studies reanalysing the RBCT data support the conclusions that culling is associated with reduced TB incidence in cattle (Mills et al. 2024a, b)

In England, a large-scale Badger Control Policy (BCP) has been in place since 2013. Under this policy badger culling is carried out over large areas (>100km^2^) under licence, with culling activities organised and implemented by members of the local farming community. In addition to badger culling, farmers within BCP areas also receive biosecurity advice and (as of 2017) additional blood testing of cattle is conducted in BCP areas once two years of badger culling have been implemented. Since 2013, a total of 70 BCP areas have been licenced across parts of the High-Risk Area (HRA) and Edge Areas of England, where TB incidence in cattle is high. Badger culling is also implemented in some parts of the Low-Risk Area, which covers Northern and Eastern England, where badgers are identified as infected with *M. bovis*.

Analyses of data from the first two BCP areas licenced in England in Gloucestershire and Somerset estimated that badger culling was associated with significant reductions in cattle TB incidence, relative to matched unculled comparison areas (Brunton et al. 2017, Downs et al. 2019). More recently, Birch et al. (2024) analysed data from 52 BCP areas using a difference in differences approach, which compared TB incidence within and between BCP areas across different time periods, both prior to and during years when BCP controls were applied. This study found further evidence that that the BCP was associated with changes in TB incidence in cattle, with an estimated average 56% reduction reached in the fourth year of BCP controls. The authors of this study acknowledge that it is difficult to separate the effects of badger culling from other disease control measures under this policy, although significant reductions in TB incidence were observed within BCP areas prior to likely effects from additional blood testing of cattle.

The results from Downs et al. (2019) and Birch et al. (2024), in combination with data from earlier culling trials in England (Donnelly et al. 2007, Jenkins et al. 2010) and Ireland (Griffin et al. 2005) offer evidence that badger culling can act to reduce levels of disease in cattle. However, these results sit in contrast to those of another recent study by Langton et al. (2022a), which failed to find evidence of any disease control benefits in cattle associated with the current badger culling policy in England. The primary analysis of this study compared culled and unculled areas within the High-Risk Area in England, but found no significant differences in cattle TB incidence rates. This study by Langton et al. (2022a) was criticised by Middlemiss and Henderson (2022) who suggested that the analysis had been carried out “in a manner that masks the effect of culling by incorrectly grouping data, which makes it impossible to assess the impact of culling on cattle TB breakdowns”. This was in turn rejected in a rebuttal by Langton et al. (2022b), which was also critical of an alternative analysis presented by the authors. Issues with reproducibility or replicability are known to occur in many fields of science and are not entirely surprising, particularly where there are differences in the analyses or methods applied by different studies (National Academies of Sciences, Engineering, and Medicine, 2019). Nevertheless, the fact that the results of Langton et al. (2022a) study suggest no impact of badger control on cattle TB incidence is a cause for concern.

The aim of this study is to investigate the robustness of the primary analyses undertaken by Langton et al. (2022a), in order to better understand how this study sits within the broader evidence base relating to badger control. Firstly, I review aspects of the data and analyses used in this study and consider how they could explain the apparent lack of effect detected. I then carry out a Monte Carlo analysis using simulated badger control data to ascertain whether the analyses used by Langton et al. (2022a) are robust enough to reliably detect possible culling effects under a range of different effect size scenarios.

### Overview of the Langton et al (2022) analyses

The primary analysis in Langton et al. (2022a) compares the cattle TB incidence in the BCP areas in the High Risk Area (HRA) of South West England with the TB incidence in unculled land in the HRA. Unlike other studies, which analysed data from multiple specific badger control areas (Downs et al. 2019, Birch et al. 2024), Langton et al. (2022a) aggregated data from multiple BCP areas in each year into a single data point, and then compared these values to the aggregated value for all herds not within BCP areas in that year. The data used by Langton et al. (2022a) is displayed in Table 1 and covers the period from 2009 to 2020. For their primary analysis the authors combine all data together into a total of 16 data points, ten data points covering the period 2009-2018 for the unculled parts of the HRA, and six data points covering the culled part of the HRA from 2013-2018.

**Table 1.** Data used in the Langton analyses, key variables are number of herds (herds), herd density (dens), number of OTF-W incidents (pos), year, culled vs unculled land (cull), time at risk (timeatrisk), TB incidents per time at risk (incpertimerisk), annual change in incidence (changeinc).

| *herd* | *dens* | *pos* | *year* | *cull* | *obs* | *timeatrisk* | *incpertimerisk* | *changeinc* | *time* |
| --- | --- | --- | --- | --- | --- | --- | --- | --- | --- |
| 24603 | NA | 2083 | 2009 | n | 1 | 19947 | 0.104 | NA | 1 |
| 23930 | NA | 2329 | 2010 | n | 2 | 17719 | 0.131 | 0.0270 | 2 |
| 22828 | NA | 2404 | 2011 | n | 3 | 17351 | 0.139 | 0.0071 | 3 |
| 22595 | NA | 2488 | 2012 | n | 4 | 16688 | 0.149 | 0.0105 | 4 |
| 22003 | 60.39 | 2367 | 2013 | n | 11 | 16161 | 0.146 | -0.0026 | 5 |
| 21369 | 58.65 | 2496 | 2014 | n | 12 | 16050 | 0.156 | 0.0091 | 6 |
| 20292 | 56.04 | 2207 | 2015 | n | 13 | 15800 | 0.140 | -0.0158 | 7 |
| 18181 | 54.77 | 1927 | 2016 | n | 14 | 13466 | 0.143 | 0.0034 | 8 |
| 15467 | 53.80 | 1485 | 2017 | n | 15 | 11241 | 0.132 | -0.0110 | 9 |
| 11144 | 48.22 | 942 | 2018 | n | 16 | 8298 | 0.114 | -0.0186 | 10 |
| 372 | 65.61 | 46 | 2013 | y | 5 | 274 | 0.168 | 0.0188 | 5 |
| 368 | 64.90 | 41 | 2014 | y | 6 | 274 | 0.150 | -0.0183 | 6 |
| 503 | 63.67 | 60 | 2015 | y | 7 | 397 | 0.151 | 0.0015 | 7 |
| 2524 | 66.37 | 295 | 2016 | y | 8 | 1834 | 0.161 | 0.0097 | 8 |
| 5159 | 62.51 | 492 | 2017 | y | 9 | 3682 | 0.134 | -0.0272 | 9 |
| 9116 | 65.63 | 771 | 2018 | y | 10 | 6520 | 0.118 | -0.0154 | 10 |

The authors use a series of Generalised linear models (GLMs), generalised linear mixed models (GLMMs), generalised additive models (GAMs) and generalised additive mixed models (GAMMs) to compare the incidence and prevalence of new breakdowns in culled and unculled areas. One potential limitation as highlighted by Middlemiss and Henderson (2022) is that the culled group each year contains a value derived from a mixture of areas, some of which have been culled for only one or two years.

Analyses of data from the RBCT found that any impacts of culling on TB in cattle were not readily observed until three years of culling were completed (Jenkins et al. 2010). More recent analyses from the current Badger Control Policy did find evidence of impacts after two years, in two of three areas analysed (mean estimates −21% Somerset, −16% Gloucester), but larger impacts after four years (−37% Somerset, −66% Gloucester) (Downs et al. 2019). Similarly Birch et al. (2024) estimated reductions in TB incidence of 9.7% (95%CI 2.8-16.4%) in the first year of BCP, compared to 56% (95%CI 29.7-75.8%) in the fourth year of BCP.

Table 2 shows the six cull years (2013-2018) analysed in the Langton et al. analyses, with each year combined into a single value/data point in Table 1. The first three years of control (2013-2015) consist of a small number of cull areas (two to three) which are in the early years of the process.

**Table 2.**
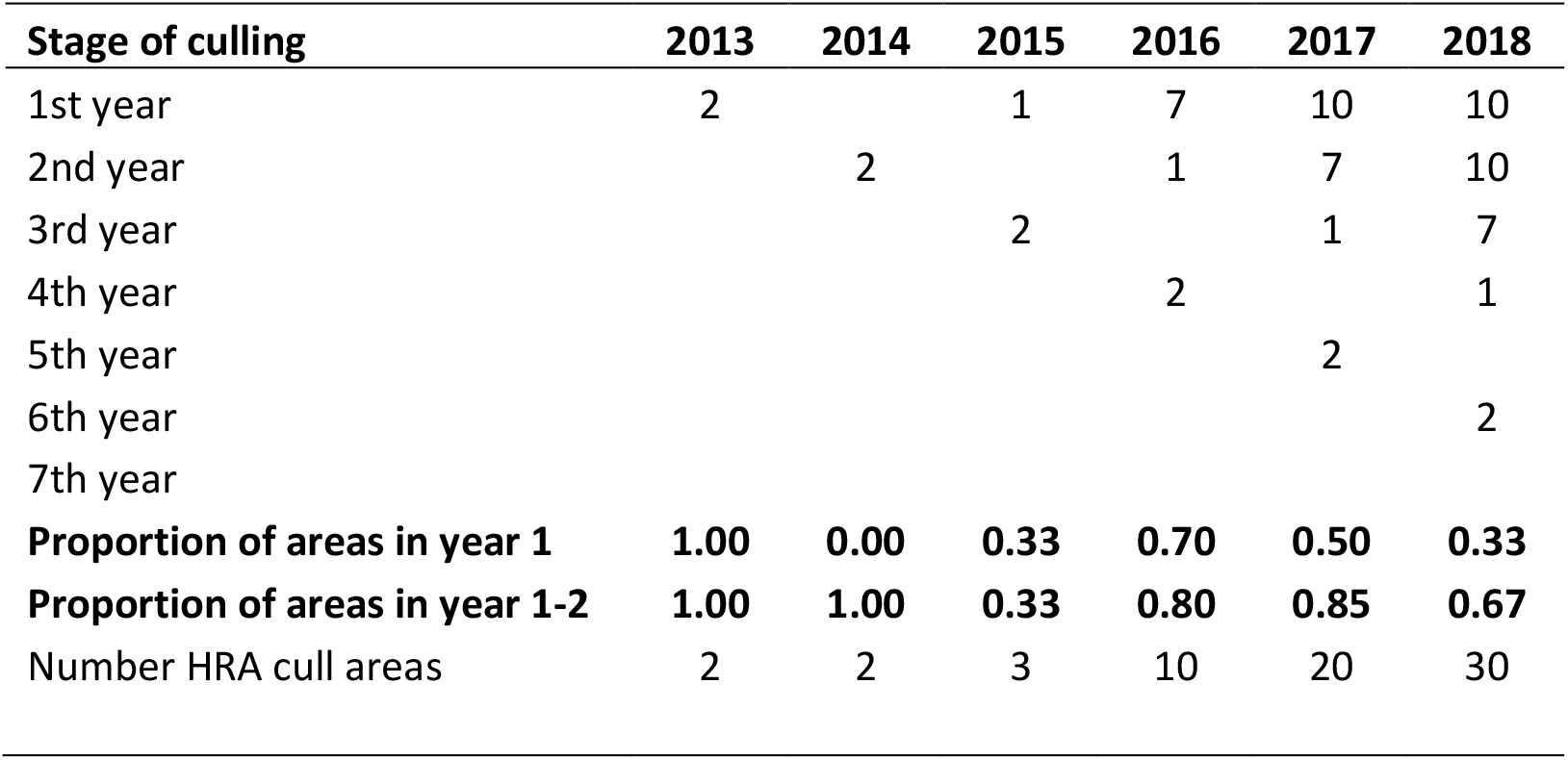
Number of HRA cull areas by the stage of culling during the period 2013-2018 (analysed by Langton et al.)

| Stage of culling | 2013 | 2014 | 2015 | 2016 | 2017 | 2018 |
| --- | --- | --- | --- | --- | --- | --- |
| 1st year | 2 |  | 1 | 7 | 10 | 10 |
| 2nd year |  | 2 |  | 1 | 7 | 10 |
| 3rd year |  |  | 2 |  | 1 | 7 |
| 4th year |  |  |  | 2 |  | 1 |
| 5th year |  |  |  |  | 2 |  |
| 6th year |  |  |  |  |  | 2 |
| 7th year |  |  |  |  |  |  |
| **Proportion of areas in year 1** | **1.00** | **0.00** | **0.33** | **0.70** | **0.50** | **0.33** |
| **Proportion of areas in year 1-2** | **1.00** | **1.00** | **0.33** | **0.80** | **0.85** | **0.67** |
| Number HRA cull areas | 2 | 2 | 3 | 10 | 20 | 30 |

Although the number of cull areas increases significantly in 2016, the vast majority of BCP areas in this sample (70%) are in year one of the process. The combined TB incidence value for 2016 (Table 1), therefore, mainly consists of herds where impacts of culling are likely to be low. The number of cull areas further increases in 2017, but 85% of areas are in the first two years of culling where impacts are likely to be low. The year 2018 is the only year with substantial numbers of areas (n=10) in the later years of the culling process (years >2), but even in this year, two thirds of areas (67%) are in the first two years of the process.

In addition to time since the intervention started, there are multiple other factors which influence TB risk in cattle, including herd size, herd type, cattle movements, previous TB history, as well as wildlife related factors (Broughan et al. 2016, McKinley et al. 2018). Previous studies investigating the impacts of badger culling accounted for these effects by following the same cattle populations over time and by controlling for these factors in multivariate analyses (Jenkins et al. 2010, Downs et al. 2019), and in the case of the RBCT, conducting a large replicated randomised trial (Bourne et al. 2007, Jenkins et al. 2010) where culling areas were matched to unculled controls.

When comparing two different populations of cattle it is possible that they differ in their TB risk, not only due to the control measures applied, but also due to the influence of these other factors. For example, in the Langton et al. data set, in 2015 the TB incidence rate is derived from 503 herds in the culled treatment and from 20,292 herds in the unculled treatment. If these two groups of herds differ in their average herd type, size or TB history, then these two groups may have different TB incidence rates, regardless of the level or impact of badger control conducted.

Evaluating the effect of any intervention or treatment may be complicated by the presence of selection biases, whereby the treatment of interest is applied in a non-random fashion. If there are consistent differences between culled and unculled areas this could also result in a degree of selection bias in the data, for example if culling is applied to areas with higher baseline incidence rates. While the Langton et al. study includes herd density in some analyses, other factors, such as prior TB history, herd type or size are not considered and it is assumed that there is no selection bias in how culling is applied.

I used freely available public data for TB incidence rates in individual BCP areas (APHA 2024) and data on TB incidence rates in the wider HRA in England (DEFRA 2025) to investigate whether there was evidence of an underlying selection bias in the application of badger control. I compared Official TB Free Withdrawn (OTF-W, where *M. bovis* infection is confirmed by culture) incidence rates for herds in existence in the calendar year that each BCP area started, to the OTF-W incidence rate in the whole HRA and to the unculled part of the HRA (Figure 1). Data from the first 30 BCP areas licensed for culling in the HRA (2013 – 2018) were used for this comparison, to cover the period analysed by Langton et al. (2022a). Inspection of the data shows a clear pattern, with values for BCP areas (the points) generally higher than both the whole HRA, or the unculled part of the HRA (Figure 1). This was further confirmed using a paired T-test which showed that the difference is statistically significant when comparing BCP areas to the whole HRA (*t*_29_=3.70, p=0.0001, mean difference = 3.74, CI=1.67-5.80), or to the unculled part of the HRA (*t*_29_=4.32, p=0.0002, mean difference = 4.35, CI=2.29-6.42). This indicates that badger culling was more likely to be applied to areas with high TB incidence rates, which could act to mask any effects of culling. This pattern was not identified or considered by the Langton et al. (2022a) study.

**Figure 1.**
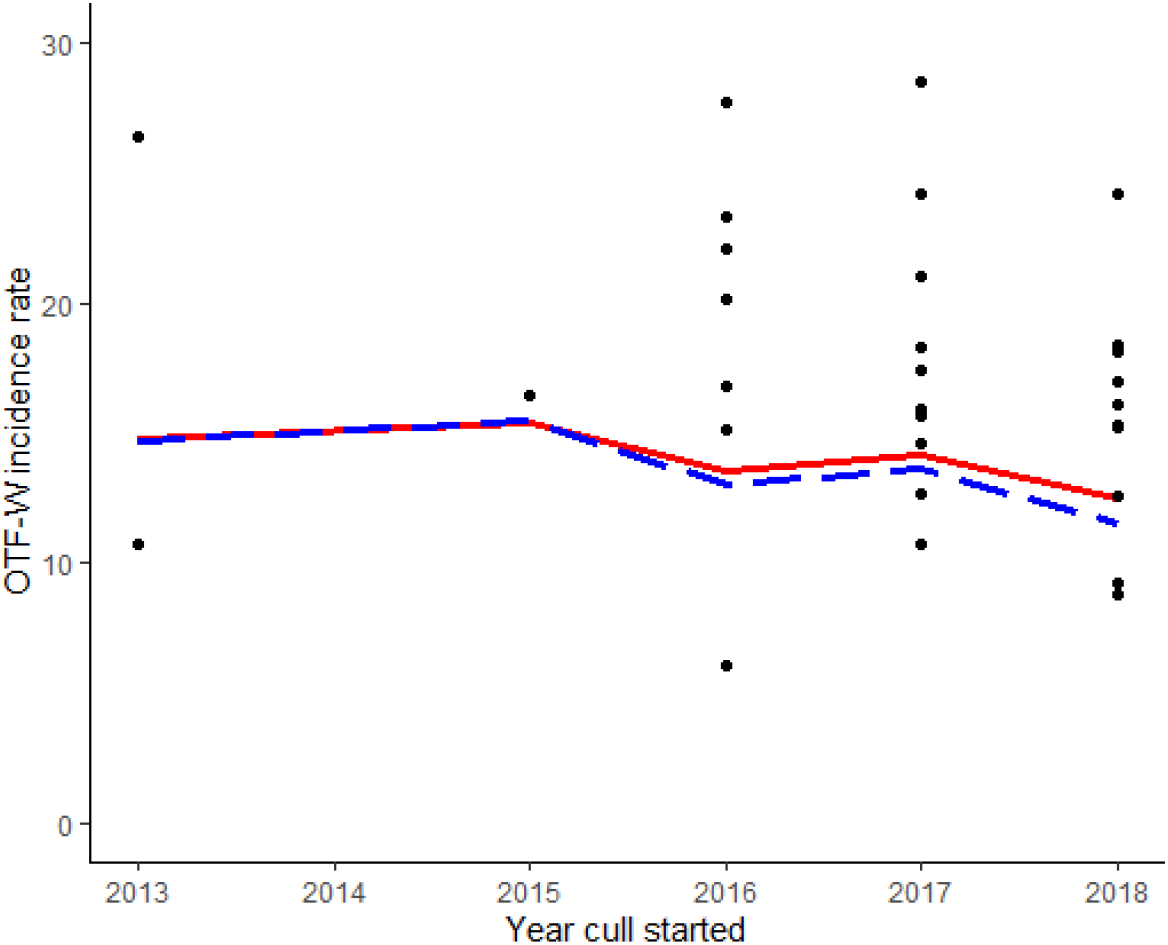
OTF-W incidence rates for herds in the BCP areas in the year badger culling started (black points). The red line is the incidence rate for the total HRA, while the blue dashed line is the incidence rate for the unculled part of the HRA. The fact that the data points are consistently above the line (rather than spread either side of it) visually illustrates how cull areas typically have much higher incidence rates than that of the HRA as a whole.

## Monte Carlo analyses - Methods

The aim of this work was to create a simulated badger cull data set, consisting of several areas, some of which are culled and others which are not. A pre-specified effect of culling could then be applied to the culled areas in the simulation. The data can then be grouped together and analysed following the Langton et al. (2022a) approach to investigate the likelihood that these statistical methods can correctly detect the effect of badger culling specified in the simulation.

To create the simulated data set we carried out the following steps:

1. A list of 70 areas was created to cover the time period 2009-2018 (10 years of observations for each area). A total of 70 areas was chosen as the mean badger cull area size is approximately 460km^2^ meaning that the HRA in England contains approximately 70 potential areas of this size where culling could be applied.
2. To generate variability among areas, each area was given a starting TB incidence rate for the year 2012 (prior to the cull policy starting) using a random value taken from a normal distribution with mean=15.0 and sd=6.06. These parameters were calculated from the 57 BCP areas in the High Risk Area in England, using the OTF-W (Official TB Free Withdrawn) incidence rate for herds in existence in the calendar year prior to culling (APHA 2024).
3. Variation in TB incidence rate over time (in the absence of culling) was achieved by adjusting the starting TB incidence rate by a random value each year taken from a normal distribution with mean=0, sd=0.2 (whereby a 0 value would equal no change, a value of 0.2 a 20% increase, and −0.2 a 20% reduction). These values were chosen as they closely matched the annual variation in real data from culled areas prior to culling. This produced a baseline data set where areas had varying level of TB, which in turn varied over the period in question (Figure 2).
4. Badger culling was then applied to thirty of these areas following the pattern observed in the current culling programme over this period: two areas starting in 2013; one area in 2015; seven areas in 2016; ten areas in 2017; and ten areas in 2018.
5. Due to the significant difference observed between baseline incidence rates in culled areas and the rest of the HRA (Figure 1), areas were selected to start culling in a non-random fashion with a bias towards areas with a higher starting TB incidence. To achieve this, areas were ranked according to their starting TB incidence and then 30 areas were selected at random, but with a 0.9 probability they would be taken from the top 35 (highest TB) areas and a 0.1 probability they would be taken from the bottom 35 (lowest TB) areas. This generated a degree of selection bias that varied in each run of the simulation, but was broadly in line with that observed in the real data (Figure 1). Following the selection of the 30 areas to be culled, the TB incidence rates in these areas in the year prior to culling starting were then compared to the unculled HRA in that year using a paired t-test (mirroring the analyses described above in Figure 1) and the estimate recorded as a measure of *cull selection bias* in that run of the simulation (Figure 3).
6. During culling years, the TB incidence rate in all areas allocated to the culling treatment was modified to reflect possible effects (i.e., magnitude of reduction) of badger culling on TB incidence in cattle. Three potential scenarios were simulated; 1) no effect of badger culling; 2) RBCT culling effects based on Jenkins et al. (2010); and 3) BCP culling effects based on Birch et al. (2024) (Table 3). To incorporate uncertainty in the magnitude of culling effects each year, a random value was generated for the impact of culling in that year for each area, based on a uniform distribution with minimum and maximum values matching the upper and lower confidence intervals. For example, using the RBCT culling effects scenario, for cull areas in the third year of culling, the TB incidence rate was calculated as the TB incidence rate in the year prior to culling, adjusted by a random value between −57.6% and −13.4% (table 3). Figure 4 below shows an example of the impact of culling in the first 10 cull areas (started in 2013-2016) using culling scenario two.
7. Once culling was applied to all areas, the cattle TB incidence rate data could then be grouped following the approach in Langton et al. (2022a). In each year 2009 to 2018, all culled areas were combined to produce a single cattle TB incidence rate. Likewise, all non-culled areas in each year were summed together to produce a single TB incidence rate. These data were compiled in a summary table matching the data in the supplementary materials from Langton et al. (2022a) (as in Table 1). TB incidence rates were converted to number of incidents assuming all simulated areas had the same time at risk (200 herd years at risk – which is broadly comparable to average BCP area).
8. Several linear models were then applied to the data, following the R code in the Langton et al. (2022a) supplementary materials. In all models the response variable was the number of TB incidents (Poisson distribution) and included an offset variable. The models applied were (*with models in R syntax*):
  a. Model 1: A General Linear Model (GLM) comparing the number of TB incidents in culled and unculled treatments with formula: *glm(pos~cull+offset(log(risk)), family=“poisson”)*
  b. Model 2: A General Additive Mixed Model (GAMM) with a linear time covariate, random effect of year and formula: *gamm4(pos~cull+time+offset(log(risk)), random=~(1*|*year), family=“poisson”)*
  c. Model 3:A General Additive Mixed Model (GAMM) with a non-linear time covariate, random effect of year and formula: *gamm4(pos~cull+t2(time)+offset(log(risk)), random=~(1*|*year), family=“poisson”)*
  In all models above *pos*=number of TB incidents, *cull*= either culled or unculled treatment, and *risk*= time at risk (used as an offset term to control for differing time at risk values in each year).
9. For comparative purposes and to validate that culling effects were adequately applied to the simulation, we also carried out an additional area level GLM. This model used the individual data for each culled area over time, where the response was the number of TB incidents in each area in each year. Fixed effects included a categorical fixed effect for area id and a categorical effect for culling stage (six levels: pre-cull, year one, year two, year three, year four and years four+). A continuous measure of time (calendar year) was also included. This model (Model 4) has the following R formula: *glm(pos ~area+year+cull_stage, family=“poisson”)*. This model does not need a time at risk variable, as all areas in the simulation have the same value.
10. For each run of the simulation the significance of these models was recorded (against a p value of 0.05) and the effect size of the cull variable was also recorded.

**Figure 2.**
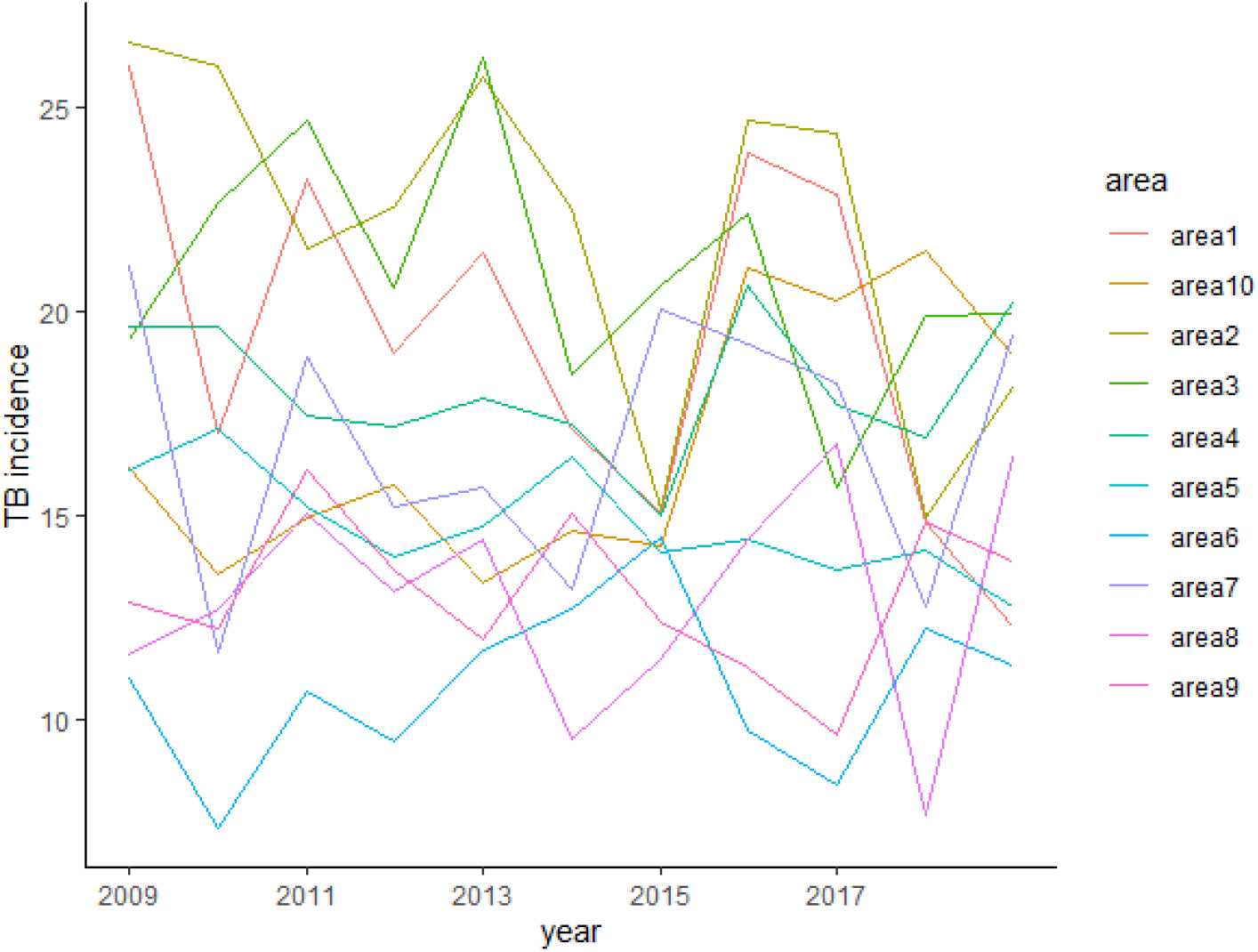
Example data from the simulation, displaying 10 (of 70) areas with varying TB incidence rate, prior to the application of culling effects.

**Figure 3.**
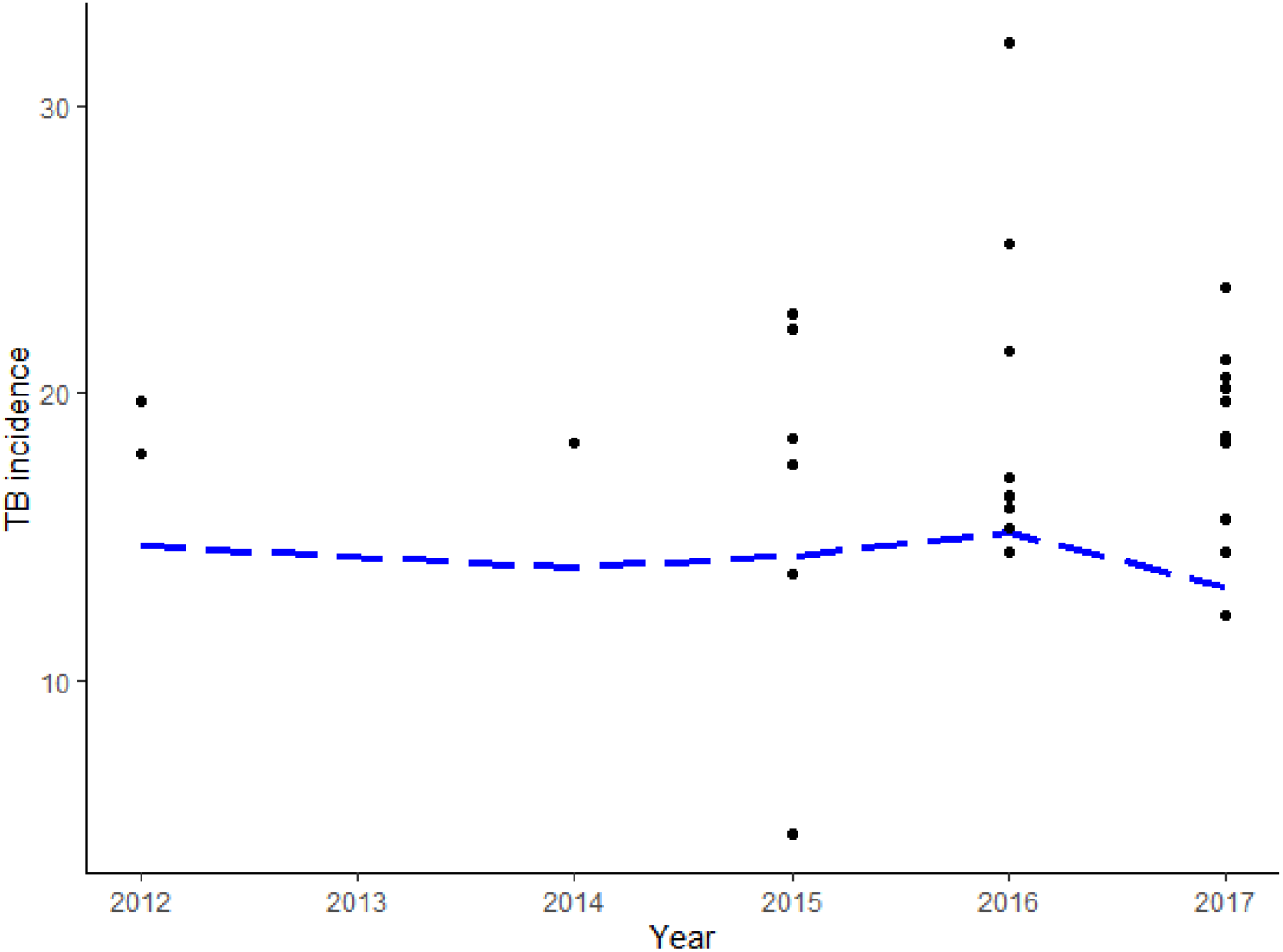
Simulated data showing an example of cull selection bias replicating patterns observed in the real data (Figure 2). Points are the TB incidence rates of individual simulated areas allocated to the culling treatment in the year prior to culling commencing (e.g data displayed for 2012 is for areas commencing culling in 2013). The blue line is the combined TB incidence rate in unculled areas in that year. In this example the cull selection bias (estimated from paired t-test) is 4.15.

**Table 3.**
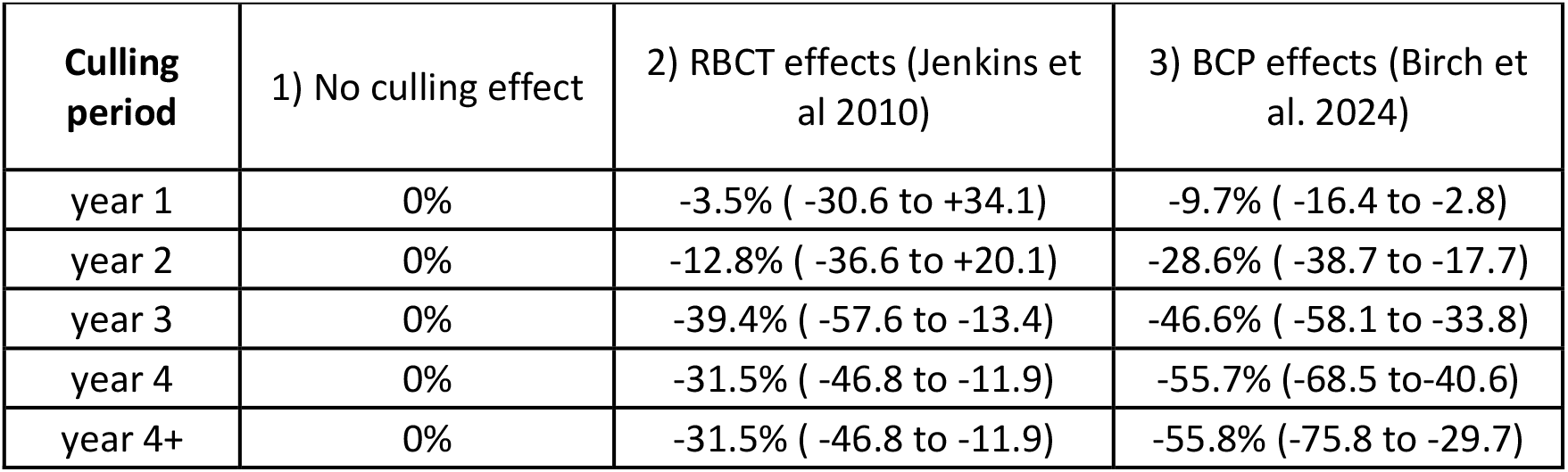
The three scenarios simulated and the culling effects (with 95% CI) in each time period.

**Figure 4.**
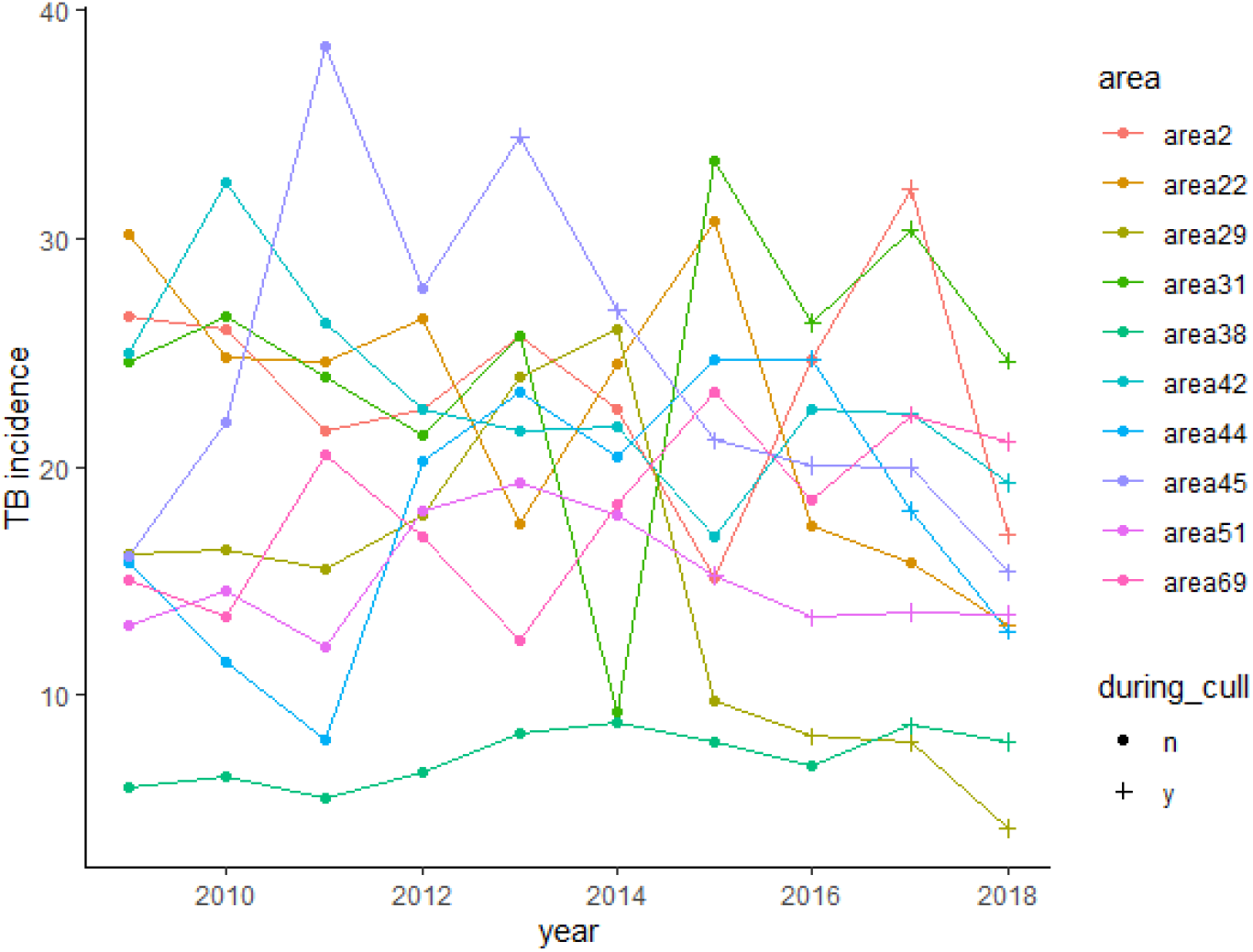
Example simulated culling effect data simulated according to the observed magnitudes of RBCT culling effects

The above steps were repeated 1000 times for each of the three culling effect scenarios and the results were combined in a summary table. This was then used to calculate the % of simulation runs where a statistically significant effect of culling was recorded, and the direction of this effect. This gives an estimate of the probability that the Langton et al. statistical methods will find an effect of culling in a simulated data set, where the effect of culling is pre-specified and known, and where TB incidence rate varies among locations and years.

## Results

The performance of the statistical models, measured as the percentage of simulations for which the model detected a significant effect of culling, depended on the magnitude of the cull effect simulated and the type of statistical model used (Table 4).

**Table 4.**
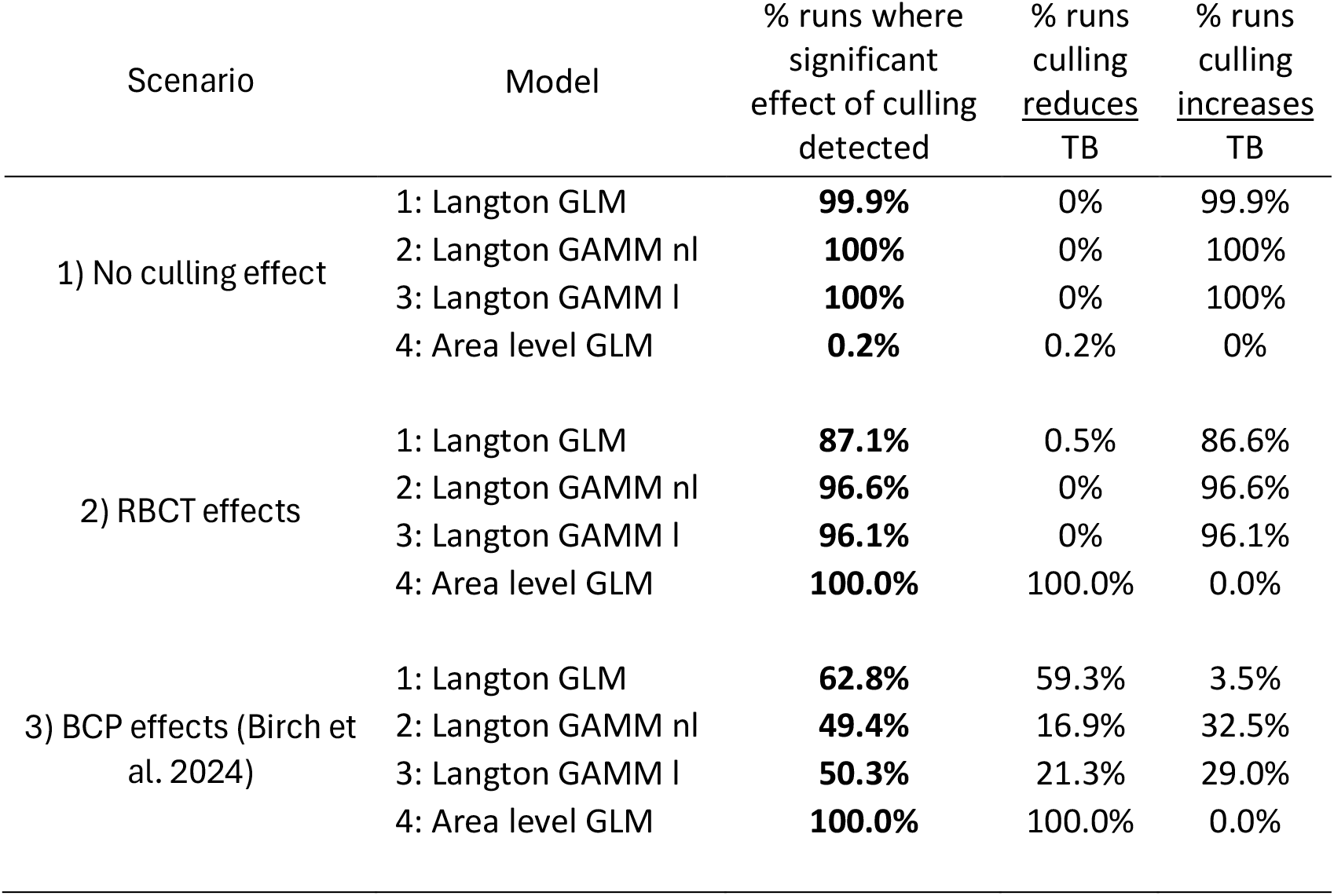
Performance of the statistical models to detect simulated culling effects under three different culling effect scenarios. For each type of statistical model, the percentage of simulation runs is displayed where a significant statistical effect of culling was detected (i.e. a difference between the culled and unculled treatments, p=<0.05). Also displayed is the percentage of runs where the model estimated that culling reduced TB incidence rate (negative coefficient in the model, culled areas lower TB incidence rate than unculled), and the percentage where culling increased TB incidence rate (positive coefficient in the model, culled areas higher TB incidence rate than unculled).

In the simulations with no effect of culling, areas were allocated to the cull treatment group, but no actual cull effect was applied in the simulation. Despite this absence of cull effect, the analyses using models 1-3 from the Langton et al. (2022) study consistently found that the cull treatment group had a higher TB incidence rate than the unculled treatment (Figure 5, Table 4). This likely reflects the selection bias in the simulation, whereby higher TB incidence areas were disproportionately (but realistically) allocated to the cull treatment. In contrast, the area level GLM (model 4) which controlled for differences among areas only found a significant effect of culling in 0.2% of simulations (Table 4).

**Figure 5.**
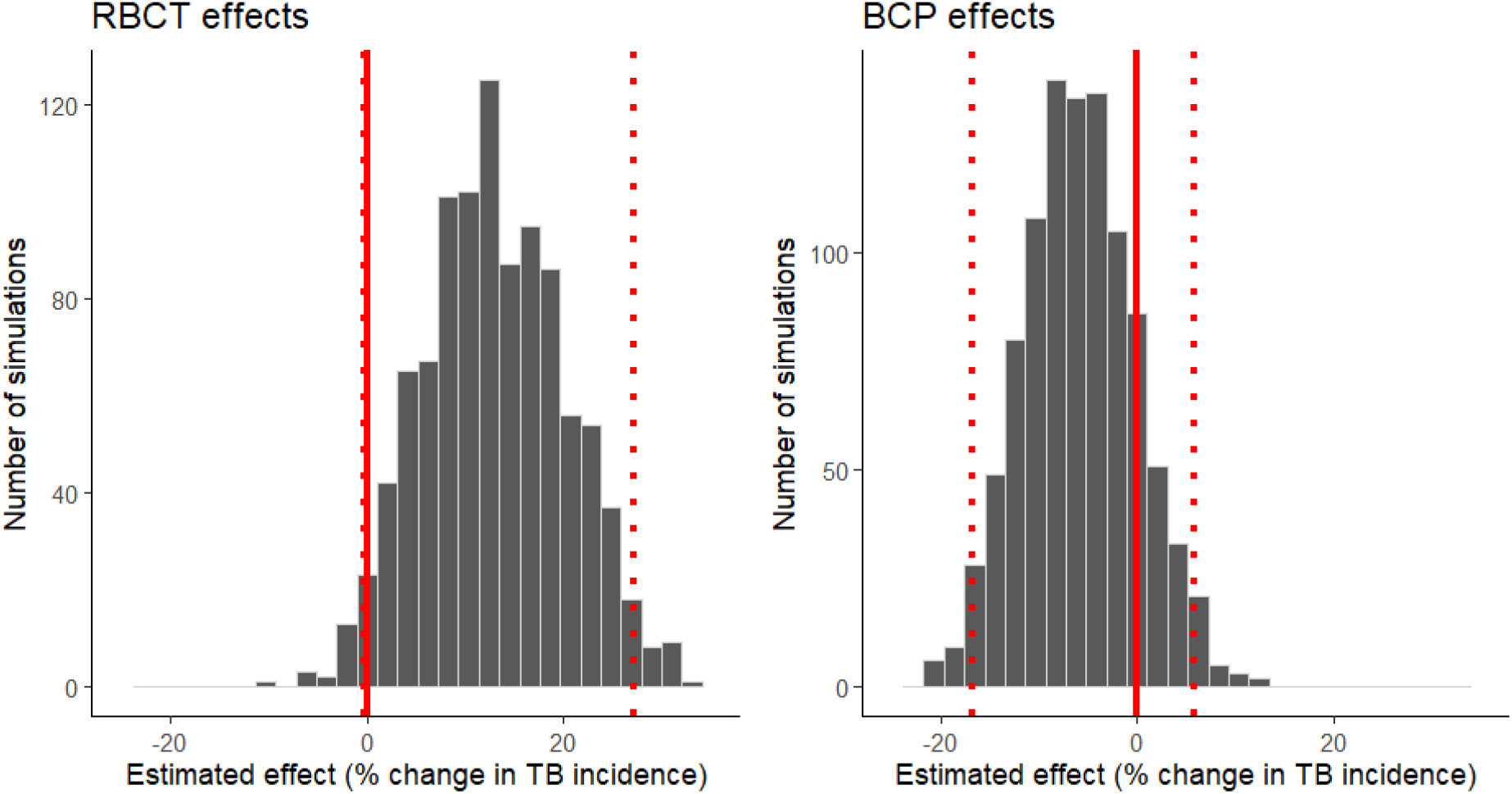
Estimated effect of culling (coefficient estimating the difference between culled and unculled areas) from GLM analyses following Langton et al. (Model 1) when applied to simulated data with cull effects. The histograms display the distribution of effect sizes estimated across 1000 simulations where (a) RBCT effect sizes were simulated (Jenkins et al 2010) and (b) BCP effect sizes were simulated (Birch et al 2024). The red line is at 0 and the dashed lines represent 95% of the data.

In the simulations with RBCT level culling effects the area level GLM correctly identified that culling resulted in a significant reduction in TB incidence in 100% of simulations (Table 4), with the estimated effects in the different cull stages (Figure 6) closely matching those applied in the simulation (Table 3). In contrast, models 1-3 again consistently estimated that culling increased TB incidence rate (Figure 5) in more than 80% of simulations, with only a small number of model 1 simulations correctly identifying a that culling was associated with a decrease in TB incidence.

**Figure 6.**
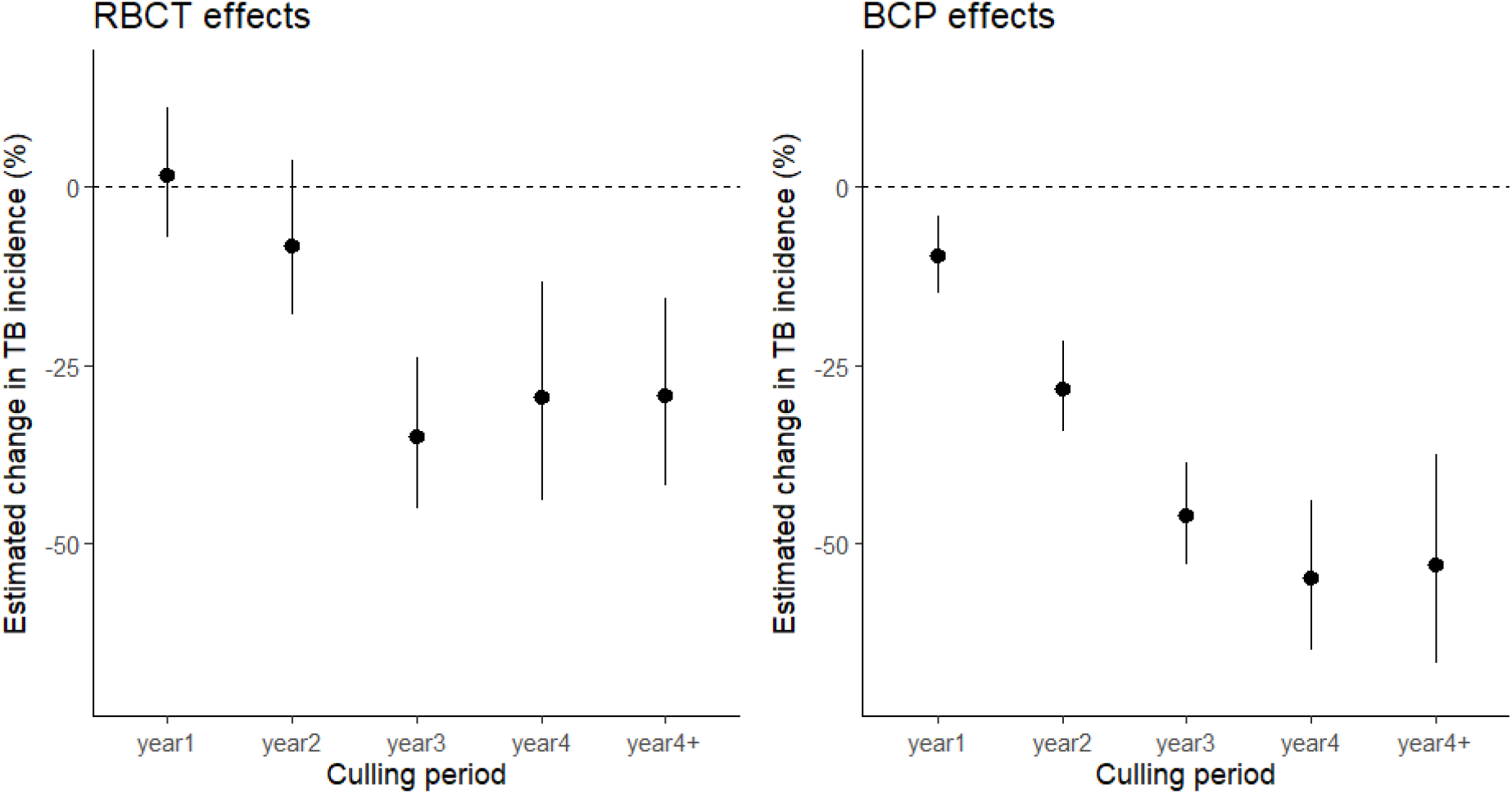
Estimated effect of culling from area level GLM analyses (Model 4), conducted on simulated data with RBCT culling effects and BCP culling effects (Table 3). Points are mean coefficients, with lines representing 95% of the estimates across all 1000 simulations.

In the simulations with BCP effects applied, the impact of culling was much greater in the early years of the cull process (Table 3). Again, the area level GLM (model 4) consistently identified that culling was associated with a significant reduction in TB incidence rate with estimated effects for specific cull stages (Figure 6) matching those specified in the simulation (Table 3). Grouping the cull areas together and comparing the TB incidence rate in this aggregated cull treatment group to the unculled area using Langton et al. models 1-3 did correctly identify a benefit of culling (reduction in cattle TB incidence rate) in some simulations with the BCP effects (Table 4). A significant benefit of culling was correctly identified in 59.3% of simulations using model 1 (Langton GLM, Figure 5). The results using the GAMM models (model 2-3), which contain a covariate for time, were less consistent; they correctly identified a significant reduction in TB incidence rate due to culling in 16.9% and 21.3% of simulations for model 2 (GAMM with nonlinear time covariate) and model 3 (GAMM with linear time covariate), respectively.

Cull selection bias was calculated for each simulation using the mean difference in TB incidence rate between areas selected for culling in a given year and unculled land in the same year using a paired t-test. Cull selection bias varied among simulation runs, due to the random nature of the simulation parameters. For the 1000 simulations with the BCP culling effects the mean cull selection bias was 3.97, with a standard deviation of 1.02, which was similar to the bias in other no cull effect (mean=3.97, sd=1.07) and RBCT effect scenarios (mean=3.94, sd=1.08). The likelihood that cull effects were correctly identified was significantly correlated with the level of selection bias in the simulation (Figure 7, r_998_=0.9, p=<0.001). Simulation runs where areas selected for culling had higher baseline TB incidence rate relative to unculled areas (a higher cull selection bias) were less likely to result in models 1-3 correctly identifying a benefit of culling. Simulation runs with only a moderate selection bias, such that cull areas only had slightly higher baseline incidence rate were more likely to correctly find a beneficial effect of culling (Figure 7). Removing cull selection bias entirely from the simulation resulted in all four models (Model 1-4) correctly identifying the significant effect of culling in close to 100% of simulations.

**Figure 7.**
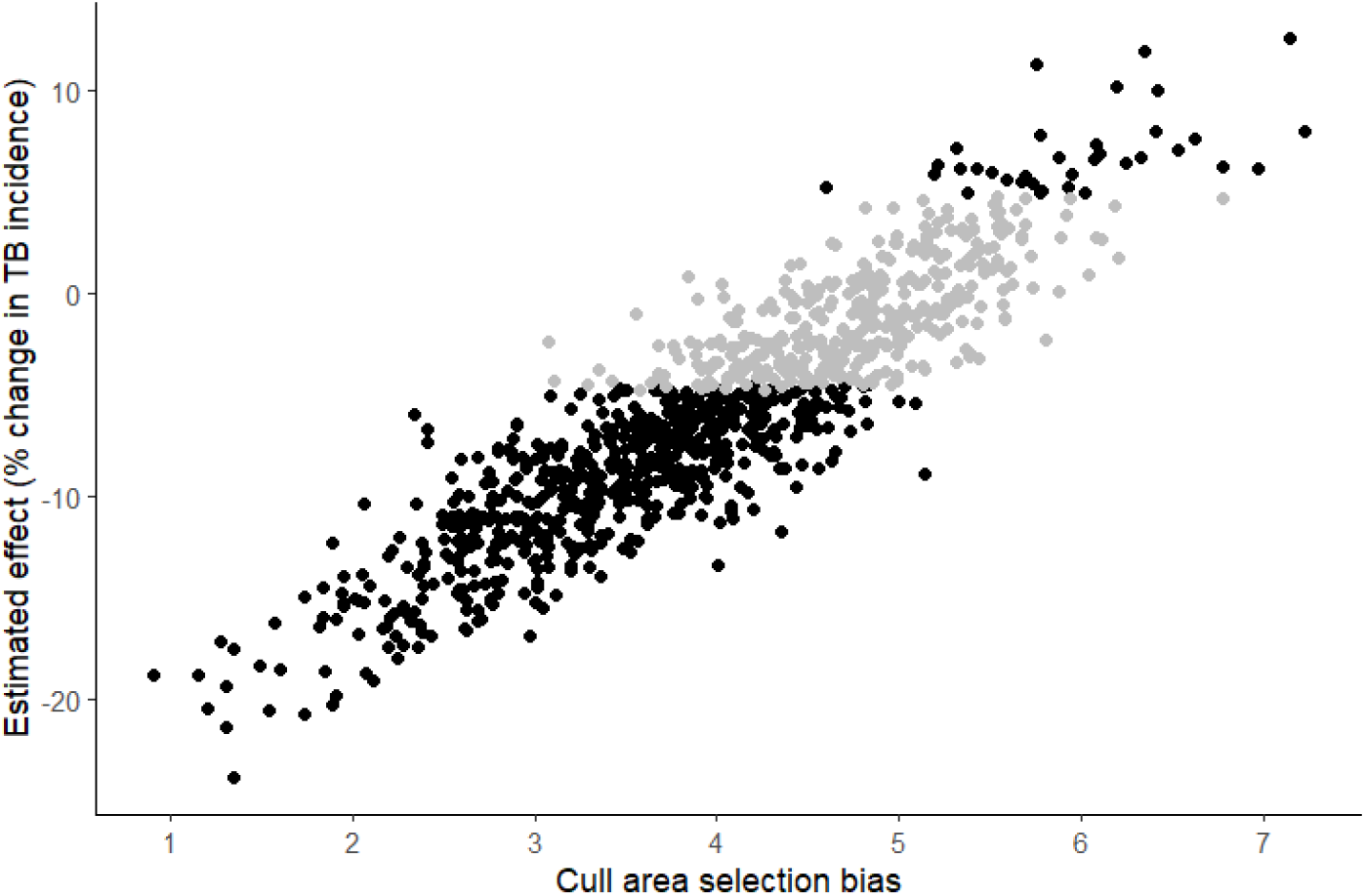
Relationship between cull selection bias (mean difference between TB incidence rate in areas selected for culling and unculled land – estimated from paired t-test, see figure 3) and the estimated effect of culling in that simulation. Data are for simulations with BCP cull effects. Black points are simulation runs with a statistically significant effect of culling identified using model 1 (Langton et al. GLM).

## Discussion

I have highlighted several reasons why the analyses by Langton et al. (2022a) may not have detected any effect of badger culling on TB incidence rate in cattle. Previous studies describing an effect of badger culling on cattle TB demonstrate that there are clear temporal patterns in disease outcomes in cattle, such that impacts are greater in latter cull years and may be small or negligible in the early stages (Donnelly et al. 2007, Jenkins et al. 2010, Birch et al. 2024). The approach used by Langton et al. (2022a) analysing data which combines all cull areas together, regardless of time since culling began risks masking the influence of intervention duration on overall effects observed. This is because any areas in the latter stages of culling, where the magnitude of any cull effect is likely to be greater and more detectable, are combined with areas in the early stage of the process, where the magnitude of any effect may be smaller and harder to detect.

A clear statistical bias in the selection of cull areas is also identified, as areas selected for badger culling had significantly higher TB incidence rates than non-culled areas in their starting year. Groups of farmers and landowners apply for a licence to cull badgers and this bias may simply reflect that groups in areas with higher rates of TB in cattle are more inclined to apply for cull licences. This pattern of high TB incidence areas being selected for the Badger Control Policy has two important implications for the analysis conducted by Langton et al. (2022a), which did not consider this source of bias. It means that in each calendar year (Table 1), TB incidence rates for the unculled area of the HRA will decline, as higher TB areas start culling and are moved into the cull treatment category. It also means that the TB incidence rate in the cull treatment data points each year will increase, as high TB areas that are in the early stages of the culling process (where any impacts are likely to be smaller in magnitude) are added to the cull treatment group and so increase its average.

The potential for these biases to influence the analyses conducted by Langton et al. (2022a) were further demonstrated by the Monte Carlo analyses conducted using simulated datasets. When either no culling effects or effects similar in magnitude to those seen in the RBCT were simulated, the three models used by Langton et al. (2022a) consistently and incorrectly indicated that badger culling led to an increase in TB incidence. In the no cull effect scenario, this increase reflects the bias in the selection of cull areas, such that higher TB incidence areas are gradually allocated to the cull treatment over time, which increases the TB incidence rate in the cull treatment group and decreases that of the unculled treatment group. In the simulations with the RBCT cull effects, reductions in TB incidence were only applied in years three onwards in line with published analyses of the RBCT (Jenkins et al. 2010). The fact that simulations with RBCT effects consistently incorrectly identified an increase in TB incidence likely reflects a combination of the cull area selection biases (more clearly apparent in the no cull effect scenario) and the fact that most areas in the culled treatment are in the early stages of culling, when any culling effects will be of lower magnitude.

The BCP effect scenario contained a larger and more rapid effect of culling, with a reduction in TB incidence rate in the first two years of the process (Birch et al. 2024). Under this scenario, a reduction in TB incidence rate due to culling was correctly detected in the majority (59%) of simulations using the simplest of the three Langton et al. (2022a) models. This result reflects that even with a selection bias towards culling in areas with a higher baseline TB incidence, a benefit of culling could be identified via a simple comparison between culled and unculled land, if the effect was large enough and appeared early in the data. However, this also highlights that there was a 40% probability that a benefit would not be detected. Furthermore, the two GAMM models containing either a linear or non-linear time effect were less likely to detect a reduction in TB incidence rate, and more likely to erroneously estimate that culling led to an increase in TB. Under the BCP effects scenario TB incidence should decrease over time within the cull treatment group from 2013-2018, as the magnitude of culling effects increase. However, TB will also decline over time within the unculled treatment group as higher TB incidence areas are gradually removed due to the selection bias. By including a covariate for time (calendar year), the GAMM models may therefore attribute cull related changes to the time variable rather than cull treatment variable, resulting in the incorrect conclusions of these models when applied to the simulated datasets.

It should be noted that, as with any simulation or model, the results of the Monte Carlo analysis were likely influenced by the parameters underpinning it. In particular, the results were heavily influenced by the cull selection bias, which specified the difference between areas selected for culling and those in the rest of the data set. The specific approach taken was chosen to produce a degree of bias comparable to that observed in the real cull data, but the random nature of the sampling and starting parameters within the simulation produced a degree of variation. The analytical approaches take from Langton et al. (2022a) were able to reliably detect an effect of culling in data from simulations with no cull section bias, or with low values of cull selection bias (<3, Figure 7). However, in simulations with higher levels of selection bias which were closer to that observed in the real data from BCP areas (4.3) the results were less reliable. This suggests that a failure to account for the magnitude of selection bias in the data may be a key reason that Langton et al. (2022a) did not identify an effect of badger control in their analyses.

Overall our results may help to explain the apparent differences in conclusions between Langton et al. (2022a) and other studies where effects of badger culling were detected (Jenkins et al. 2010, Birch et al. 2024). It may also explain why the results of the Langton et al. (2022a) study report a positive direction of the culling effect in some analyses conducted. In this study, all area level GLM analyses correctly identified the effect of culling in the simulated data regardless of the cull effect scenario parameters. These results highlight the importance of controlling for variable differences among areas or subjects when trying to correctly ascertain treatment effects, and highlights that analyses which aggregate data inappropriately risk spurious or false conclusions.

## Declaration

A proportion of this analytical work was undertaken by AR while working for Natural England. AR joined Defra in 09/23 and the manuscript and final versions of the analyses were complete there.

## References

APHA. 2024. Background information: badger control areas monitoring data up to 2023. https://www.gov.uk/government/publications/bovine-tb-in-cattle-badger-control-areas-monitoring-data-up-to-2023/background-information-badger-control-areas-monitoring-data-up-to-2023.

Birch, C. P. D., M. Bakrania, A. Prosser, D. Brown, S. M. Withenshaw, and S. H. Downs. 2024. Difference in differences analysis evaluates the effects of the badger control policy on bovine tuberculosis in England. Scientific Reports 14:4849.

Bourne, J., C. Donnelly, D. Cox, G. Gettinby, I. Morrison, and R. Woodroffe. 2007. Bovine TB: The Scientific Evidence. A Science Base for a Sustainable Policy to Control TB in Cattle. An Epidemiological Investigation into Bovine Tuberculosis. The Final Report of the Independent Scientific Group on Cattle TB., London.

Broughan, J. M., D. Maye, P. Carmody, L. A. Brunton, A. Ashton, W. Wint, N. Alexander, R. Naylor, K. Ward, A. V. Goodchild, S. Hinchliffe, R. D. Eglin, P. Upton, R. Nicholson, and G. Enticott. 2016. Farm characteristics and farmer perceptions associated with bovine tuberculosis incidents in areas of emerging endemic spread. Preventive veterinary medicine 129:88–98.

Brunton, L. A., C. A. Donnelly, H. O’Connor, A. Prosser, S. Ashfield, A. Ashton, P. Upton, A. Mitchell, A. V. Goodchild, and J. E. Parry. 2017. Assessing the effects of the first 2 years of industry-led badger culling in England on the incidence of bovine tuberculosis in cattle in 2013–2015. Ecology and Evolution.

Crispell, J., C. H. Benton, D. Balaz, N. De Maio, A. Ahkmetova, A. Allen, R. Biek, E. L. Presho, J. Dale, G. Hewinson, S. J. Lycett, J. Nunez-Garcia, R. A. Skuce, H. Trewby, D. J. Wilson, R. N. Zadoks, R. J. Delahay, and R. R. Kao. 2019. Combining genomics and epidemiology to analyse bi-directional transmission of Mycobacterium bovis in a multi-host system. eLife 8:e45833.

DEFRA. 2025. TB in cattle in Great Britain - headline statistical dataset https://www.gov.uk/government/statistical-data-sets/tuberculosis-tb-in-cattle-in-great-britain.

Donnelly, C. A., and P. Nouvellet. 2013. The Contribution of Badgers to Confirmed Tuberculosis in Cattle in High-Incidence Areas in England. PLoS Currents 5:ecurrents.outbreaks.097a904d093f3619db3612fe3678d3624bc776098.

Donnelly, C. A., G. Wei, W. T. Johnston, D. R. Cox, R. Woodroffe, F. J. Bourne, C. L. Cheeseman, R. S. Clifton-Hadley, G. Gettinby, P. Gilks, H. E. Jenkins, A. M. Le Fevre, J. P. McInerney, and W. I. Morrison. 2007. Impacts of widespread badger culling on cattle tuberculosis: concluding analyses from a large-scale field trial. International Journal of Infectious Diseases 11:300–308.

Donnelly, C. A., R. Woodroffe, D. R. Cox, F. J. Bourne, C. L. Cheeseman, R. S. Clifton-hadley, G. Wei, G. Gettinby, P. Gilks, H. Jenkins, W. T. Johnston, A. M. L. Fevre, J. P. Mcinerney, and W. I. Morrison. 2006. Positive and negative effects of widespread badger culling on tuberculosis in cattle. Nature 439:843–846.

Downs, S. H., A. Prosser, A. Ashton, S. Ashfield, L. A. Brunton, A. Brouwer, P. Upton, A. Robertson, C. A. Donnelly, and J. E. Parry. 2019. Assessing effects from four years of industry-led badger culling in England on the incidence of bovine tuberculosis in cattle, 2013–2017. Scientific Reports 9:14666.

Enticott, G. 2015. Public attitudes to badger culling to control bovine tuberculosis in rural Wales. European Journal of Wildlife Research 61:387–398.

Griffin, J., D. Williams, G. Kelly, T. Clegg, I. O’boyle, J. Collins, and S. More. 2005. The impact of badger removal on the control of tuberculosis in cattle herds in Ireland. Preventive veterinary medicine 67:237–266.

Jenkins, H. E., R. Woodroffe, and C. A. Donnelly. 2010. The duration of the effects of repeated widespread badger culling on cattle tuberculosis following the cessation of culling. PLoS ONE 5:e9090.

Langton, T. E., M. W. Jones, and I. McGill. 2022a. Analysis of the impact of badger culling on bovine tuberculosis in cattle in the high-risk area of England, 2009–2020. Veterinary Record 190:e1384.

Langton, T. E., M. W. Jones, and I. McGill. 2022b. Badger culling to control bovine TB. Veterinary Record 190:289–290.

Little, T., P. Naylor, and J. Wilesmith. 1982. Laboratory study of Mycobacterium bovis infection in badgers and calves. The Veterinary Record 111:550–557.

McKinley, T. J., D. Lipschutz-Powell, A. P. Mitchell, J. L. N. Wood, and A. J. K. Conlan. 2018. Risk factors and variations in detection of new bovine tuberculosis breakdowns via slaughterhouse surveillance in Great Britain. PLoS ONE 13:e0198760.

Medicine, N. A. o. S. E. a. 2019. Reproducibility and replicability in science. National Academies Press.

Middlemiss, C., and G. Henderson. 2022. Badger culling to control bovine TB. Veterinary Record 190:243–244.

Mills, C. L., R. Woodroffe, and C. A. Donnelly. 2024a. An extensive re-evaluation of evidence and analyses of the Randomised Badger Culling Trial (RBCT) I: Within proactive culling areas. Royal Society Open Science 11:240385.

Mills, C. L., R. Woodroffe, and C. A. Donnelly. 2024b. An extensive re-evaluation of evidence and analyses of the Randomised Badger Culling Trial II: In neighbouring areas. Royal Society Open Science 11:240386.

Palmer, M. V., T. C. Thacker, W. R. Waters, C. Gortázar, and L. A. L. Corner. 2012. Mycobacterium bovis: A Model Pathogen at the Interface of Livestock, Wildlife, and Humans. Veterinary medicine international 2012:1–17.

Torgerson, P. R., S. Hartnack, P. Rasmussen, F. Lewis, and T. E. S. Langton. 2024. Absence of effects of widespread badger culling on tuberculosis in cattle. Scientific Reports 14:16326.

Torgerson, P. R., S. Hartnack, P. Rasmussen, F. I. Lewis, P. O’Donnell, and T. E. S. Langton. 2025. Randomised Badger Culling Trial—no effects of widespread badger culling on tuberculosis in cattle: comment on Mills, Woodroffe and Donnelly (2024a, 2024b). Royal Society Open Science 12:241609.

van Tonder, A. J., M. J. Thornton, A. J. K. Conlan, K. A. Jolley, L. Goolding, A. P. Mitchell, J. Dale, E. Palkopoulou, P. J. Hogarth, R. G. Hewinson, J. L. N. Wood, and J. Parkhill. 2021. Inferring Mycobacterium bovis transmission between cattle and badgers using isolates from the Randomised Badger Culling Trial. PLoS Pathogens 17:e1010075.

